# PolNet Analysis: a software tool for the quantification of network-level endothelial cell polarity and blood flow during vascular remodelling

**DOI:** 10.1101/237602

**Authors:** Miguel O. Bernabeu, Martin L. Jones, Rupert W. Nash, Anna Pezzarossa, Peter V. Coveney, Holger Gerhardt, Claudio A. Franco

**Affiliations:** Centre for Medical Informatics, Usher Institute, The University of Edinburgh, 9 Little France Road, Edinburgh EH16 4UX, United Kingdom.; Centre for Computational Science, Department of Chemistry, University College London, London WC1H 0AJ, United Kingdom.; Electron Microscopy Science Technology Platform, The Francis Crick Institute, Lincolns Inn Fields Laboratories, 44 Lincolns Inn Fields, London WC2A 3LY, UK.; EPCC, School of Physics and Astronomy, The University of Edinburgh, James Clerk Maxwell Building, Peter Guthrie Tait Road, Edinburgh, EH9 3FD, United Kingdom.; Instituto de Medicina Molecular, Faculdade de Medicina, Universidade de Lisboa, Avenida Professor Egas Moniz, 1649-028, Lisboa, Portugal.; Max-Delbrück Center for Molecular Medicine, Berlin, Germany.; Vascular Patterning Laboratory, Vesalius Research Center, Leuven, Belgium.; Department of Oncology, Vascular Patterning Laboratory, Vesalius Research Center, Leuven, Belgium.; German Center for Cardiovascular Research, Berlin, Germany.; Berlin Institute of Health, Berlin, Germany.

## Abstract

In this paper, we present PolNet, an open source software tool for the study of blood flow and cell-level biological activity during vessel morphogenesis. We provide an image acquisition, segmentation, and analysis protocol to quantify endothelial cell polarity in entire *in vivo* vascular networks. In combination, we use computational fluid dynamics to characterise the haemodynamics of the vascular networks under study. The tool enables, for the first time, network-level analysis of polarity and flow for individual endothelial cells. To date, PolNet has proven invaluable for the study of endothelial cell polarisation and migration during vascular patterning, as demonstrated by our recent papers [1, 2]. Additionally, the tool can be easily extended to correlate blood flow with other experimental observations at the cellular/molecular level. We release the source code of our tool under the LGPL licence.

## 1 Introduction

The establishment of a functional, patterned vascular network is crucial for development, tissue growth, and physiology. Conversely, mispatterning of vascular networks contributes to the pathogenesis of several diseases, including arteriovenous malformations, haemangioma, hereditary haemorrhagic telangiectasias or venous cavernomas. Sprouting angiogenesis is one of the mechanisms responsible of expanding vascular networks into avascular areas. This process only generates a dense and immature network of vessels that requires subsequent extensive remodelling to form a functional, hierarchically branched network – a process termed vascular remodelling [3]. In contrast to sprouting, the molecular mechanisms regulating vascular remodelling are poorly understood. The forces exerted by blood on the luminal surface of the endothelium (most notably wall shear stress, WSS) have been recognised as a primary driver regulating vascular remodelling [4, 5, 6]. Recent work by the authors and others identified that endothelial WSS regulates endothelial cell polarity and cell migration, orchestrating vascular remodelling [7, 8, 1, 2] and controlling vessel diameter [9, 10, 11, 12]. Thus, there is an increasing interest in understanding mechanistically how haemodynamic forces impact endothelial response at the cellular and molecular level.

The mouse retina is one of the most popular models to investigate the molecular mechanisms governing angiogenesis. However, how blood flow influences the development of the retinal vasculature remained elusive. Progress is hampered by the limitations of current assays used to probe the relationship between haemodynamics and molecular response, and the complexity of *in vivo* vascular connectivity. To date, researchers have primarily used in *vitro* microfluidic assays to study the impact of blood flow on endothelial cell biology, and extrapolate these observations to explain phenotypic changes in mouse mutants. Even though some authors have been able to measure microvascular WSS *in vivo* (see Lipowsky et al. [13] for a survey), this has never been achieved, to the best of our knowledge, in the context of vascular morphogenesis in the mouse retina model.

Our recent work on vascular remodelling established a strong connection between blood flow and endothelial cell polarity. We developed PolNet to be able to quantify the relationship between endothelial cell polarity and WSS. PolNet’s image processing algorithms and computational fluid dynamics (CFD) simulator were described and validated in Bernabeu et al. [14]. PolNet was then successfully used in two recent publications: a) in Franco et al. [1], we showed that flow-induced cell polarisation directs migration of endothelial cells away from low flow segments; and b) in Franco et al. [2], we showed that non-canonical Wnt signalling modulates the endothelial shear stress flow sensor during vascular remodelling.

## 2 Methods

The design and implementation of PolNet is better understood in the context of the complete protocol, comprising experimental and computational parts, used in Franco et al. [1, 2]. Materials and setup are described in Supplementary Material Section A and a step-by-step protocol is provided in Supplementary Material Section B. Briefly, samples of murine retinal plexus are collected at different postnatal (P-) days, fixed, and labelled for luminal surface (ICAM2), cell nuclei (Erg), and Golgi apparatus (Golph4). The retinal vascular networks are imaged and post-processed to generate a binary mask from the ICAM2 channel and a second image with, at least, the Erg and Golph4 channels. These two images define the input to PolNet. An example dataset is provided in Supplementary Material Section D. Based on them, PolNet can be used to quantify the relationship between endothelial cell polarity (defined for each cell as the vector originating from the centre of mass of the nucleus and directed to the centre of mass of the Golgi complex) and the direction and magnitude of the computed traction vector (*i.e*. the product of the deviatoric stress tensor and the surface normal), which we will refer to as WSS vector for convenience. PolNet provides a graphical user interface for the user to perform the following three tasks: a) to construct a flow model from the ICAM2 mask and use the HemeLB flow solver (Mazzeo and Coveney, 2008) to estimate WSS across the whole network (as well as blood velocity, shear rate, and pressure), b) to interactively delineate the cell polarity vectors for each endothelial cell in the plexus based on the Erg-GOLPH4 image, and c) to statistically analyse the relationship between the cell polarity and WSS vectors. PolNet currently offers the following analyses:

1. Directionality table: number and ratio of endothelial cells polarised against the flow direction.
2. Polar histogram with the distribution of endothelial cell polarisation angles in relation to the blood flow direction. Each plexus can also be subdivided into regions (*e.g*. artery, vein, capillary and sprouting front) and each cell will be assigned to one of these vascular beds for further analysis. Multiple angular distributions can be subsequently compared using the Kuiper test to evaluate the likelihood that the two samples are drawn from the same underlying angular distribution.
3. Wall shear stress sensor: In this analysis, we bin the angular data according to WSS magnitude to plot the proportion of cells oriented against the flow, which we define as having an angle of 180° ± 45° with regards to the flow direction. We consider the threshold for polarisation as the WSS value leading to 60% of endothelial cells aligned against the flow direction.
4. Scalar product slope: We calculate the scalar product of the polarity and flow vectors, given by magnitude(cell polarity)*magnitude(WSS)*cos(theta), where theta is the angle between them, which combines information about the length and relative angles of the vectors. By plotting the scalar product against WSS, we can study and compare between groups the plot slope, i.e. magnitude(cell)*cos(theta). A larger negative slope corresponds to a larger polarisation effect for a given WSS, which is a surrogate measure for the sensitivity of cells to flow.

### 2.1 PolNet software repositories and contributions

In Supplementary Material Section A, we described how to install and run PolNet through the Docker platform. Docker images define a virtual software environment, such as a computer programme along with its dependencies, that can be run simply and reliably on many different machines. This approach facilitates software deployment (via the Docker Hub cloud service) for users who are only interested in the functionality described in this paper.

However, PolNet can be easily modified to perform similar analysis involving the comparison of in *silico* flow estimates and other experimental observations at the cellular/molecular level. Table 1 summarises the software repositories hosting the different components of PolNet. Users interested in expanding the functionality of PolNet should obtain the code in the PolNet main repository and follow the installation instructions for developers. This will provide access to the source MATLAB and Python scripts defining the graphical user interface and the main pre- and post-processing algorithms. Users can then adapt the code to meet the requirements of their analysis and potentially contribute the changes back to the main repository. The user can build his/her own Docker container based on the instructions provided in our PolNet Dockerfile repository. Note that in order to run the HemeLB setup tool or solver as part of the PolNet developer version, a HemeLB developer installation is required. Please refer to the HemeLB repository for installation instructions. Similarly, users can report bugs and suggest new features to the PolNet developers (and other interested users) by creating new issues on the relevant GitHub repository.

**Table 1:** Software repositories hosting the code used to build the PolNet tool

| PolNet component | Components | Software repository |
| --- | --- | --- |
| PolNet main | Graphical user interface and main pre- and post-processing algorithms | https://github.com/mobernabeu/polnet |
| PolNet Dockerfile | Docker-specific instructions to create the PolNet container | https://github.com/mobernabeu/docker-polnet |
| HemeLB | Flow model setup tool and CFD solver | https://github.com/UCL/hemelb |

### 2.2 Limitations of the method

#### 2.2.1 Capillary overlapping

We employ the maximum intensity Z-projection of a confocal microscopy image stack to segment the luminal space defining the flow domain. This implies that all the information in the Z-axis is projected onto a plane and therefore the method is only applicable to capillary beds where all the vessels can be considered coplanar in this projection. In the case of the neonatal mouse retina, this condition is fulfilled before P7, where a single non-overlapping plexus covers the outer layers of the retina. If this condition is not fulfilled, vessels overlapping or going past each other at different depths will appear connected in the plexus segmentation, which will lead to inaccuracies in the simulated haemodynamics. This situation can be easily diagnosed if the image segmentation displays branching points with four afferent/efferent segments. Thus, this protocol is not suitable for mouse retinas beyond P7 or other vascular plexuses with complex 3D organisation. This would require segmentation of 3D image stacks, an aspect not yet implemented in the current version of PolNet.

#### 2.2.2 Accuracy of the flow estimates generated

The accuracy of the WSS estimates produced by the flow solver is of paramount importance for the usefulness of PolNet. Here we discuss some potential sources of error.

a. Vessel geometry reconstruction The geometry of the flow domain has a strong influence on the computed haemodynamics. Therefore, it is critical that the process of plexus sample preparation for imaging is done with great care to avoid introducing artefacts (*e.g*. artificial vessel disconnections) and to monitor that the labelling and mounting protocol does not lead to significant shrinkage of the luminal space. Furthermore, the vessel reconstruction assumes a circular cross-section. All these potential sources of error lead to uncertainty in the flow estimates that requires quantification. The user can do this by comparing the flow results against existing experimental measurements of blood velocity in the mouse retinal vasculature (see Bernabeu et al. [14] and Table 2 for values compiled from the literature).
b. Choice of inlet and outlet boundary conditions The samples under study typically comprise only a subset of the retinal vascular plexus. These take the form of wedges with an artery that runs radially from the optic disc across the wedge and feeds capillary beds located at either side of it, which in turn are drained by veins that return in the same radial fashion to the optic disc (see Franco et al. [1,2]). We term this as the vein-artery-vein (V-A-V) configuration, although other configurations are possible (*e.g*. A-V and A-V-A). As well as imposing no-slip boundary conditions at the vessel walls, we need to specify boundary conditions at the networks inlets and outlets. In Bernabeu et al. [14], we surveyed the literature for experimental measurements of blood flow rate or pressure in the central retinal artery/vein to be used as boundary conditions. We concluded that the V-A-V configuration combined with pressure boundary conditions minimizes the modelling error appearing in the capillary beds defined between each artery-vein pair (including the sprouting front) as well as in the arteries. These are our main regions of interest. Due to the lack of such measurements in the neo-natal mouse, we used values measured in adult mice. This design decision can lead to inaccuracies in the flow estimates generated, which are difficult to quantify a priori. In Bernabeu et al. [14], we performed a sensitivity analysis of the inlet/outlet pressures used in our simulations and observed that, although velocity and WSS goes up as the pressure difference between inlets and outlets increases, the main perfusion patterns (*e.g*. areas of relative low/high flow, flow direction) remain unchanged for a wide range of inlet/outlet configurations and that the relative differences in WSS that have been shown to drive remodelling [1] are also preserved.
c. Choice of rheology model Blood is a dense suspension of deformable cells in blood plasma, leading to a complex range of rheological properties appearing at different scales. When using a CFD approach to simulate blood flow, one has to choose an appropriate approximation for this rheology. In the current work, we use the homogeneous shear-thinning rheology model proposed in [14]. This approach is a compromise between accuracy and computational tractability [15]. However, it fails to capture certain haemorheological features (*e.g*. the plasma skimming and the Fåhræus-Lindqvist effects) when applied to the simulation of blood flow in small capillaries, especially as vessel calibre gets closer to, or even smaller than, the typical red blood cell (RBC) diameter of ≈ 8*μm*.

**Table 2:** WSS values reported in the literature

| Ref. | Species, tissue | Vessel | WSS (Pa) |
| --- | --- | --- | --- |
| [16] | Cat, sartorius muscle | Arterioles | 6–14 |
|  |  | Capillaries | 1.2 |
|  |  | Venules | 0.3–1 |
| [17] | Human, various tissues | Arteries | 1–7 |
|  |  | Veins | 0.1–0.6 |
| [18] | Mouse, infrarenal aorta | Artery | $8.76 \pm 0.83$ |
| | Rat, infrarenal aorta | Artery | $7.05 \pm 0.67$ |
| | Human, infrarenal aorta | Artery | $0.48 \pm 0.03$ |

## 3 Results

In this section, we present an example of PolNet analysis performed on the data in Supplementary Material Section D of this paper. We choose a subset of a P6 mouse retinal plexus for demonstration purposes. The interested reader can refer to [14, 1, 2] for analyses based on larger plexuses. Supplementary Material Section C provides troubleshooting advice.

A P6 mouse retinal plexus was prepared, imaged, and post-processed following the instructions in Supplementary Material Section B steps 1-20. A subset of interest including arteries, veins, and capillaries was identified in the resulting images and cropped to simplify further analysis. Figure 1.a shows the first output image combining the specific stainings (Golph4, Erg, and ICAM2). Figure 1.b shows the second one with the segmentation of the ICAM2 channel.

**Figure 1:**
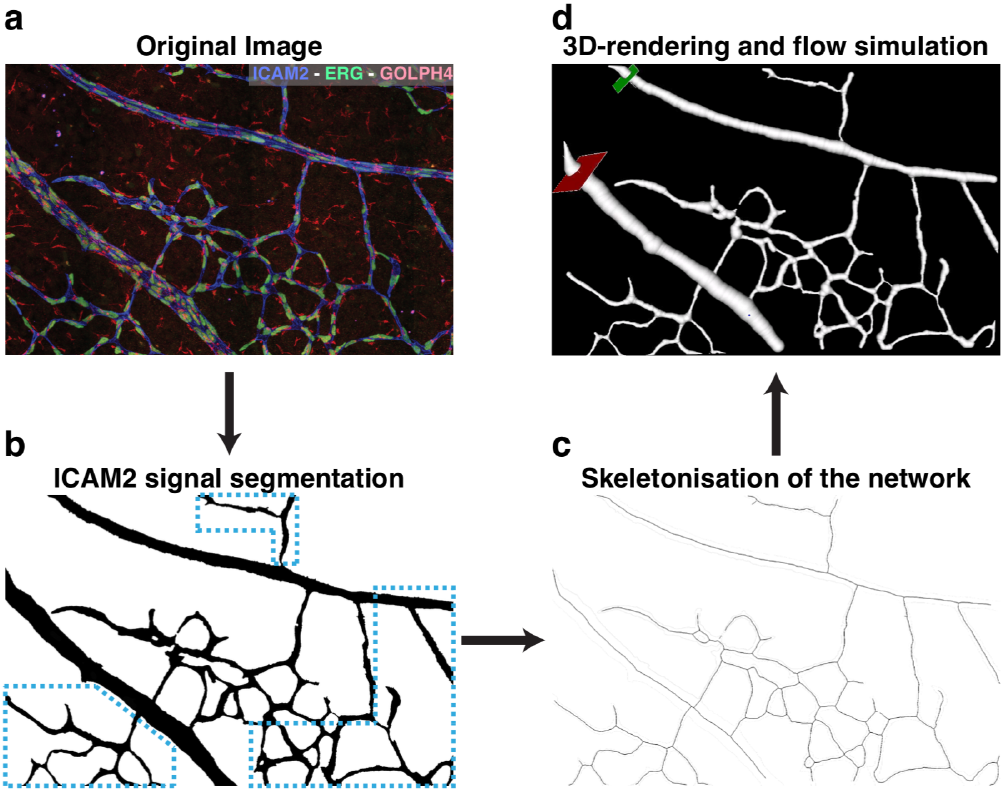
a) Example of Golph4 (red, Golgi), Erg (green, nucleus), and ICAM2 (blue, luminal surface) stainings of a subset of a P6 mouse retinal plexus. b) Binary mask generated from the ICAM2 staining in a). Note that the haemodynamics recovered in the blue highlighted region will not be accurate due to the missing connections to the nearby vessels. The interested reader can refer to [14, 1, 2] for analyses based on larger plexuses. c) Results of the skeletonisation step. d) Flow simulation setup with inlet, outlet, and seed position defined.

The ICAM2 mask was processed with PolNet in order to construct a flow model. Figure 1.c shows the results of the skeletonisation step. Figure S3.a presents the vessel calibre histogram generated by the surface reconstruction stage. Figure 1.d shows a screenshot of the flow simulation setup operation (note the location of the flow inlet in green and flow outlet in red). Finally, Figure S3.b presents the wall shear stress histogram generated at the end of the simulation. Table 2 summarises experimentally measured WSS values for simulation validation purposes.

The Golph4-Erg-ICAM2 image was used to delineate cell polarity across the whole plexus with PolNet. This is achieved by placing, on top of the image being displayed, consecutive pairs of points corresponding to the approximate centre of mass of the nucleus and Golgi of any given cell. Every time a new pair is added, an arrow connecting them is automatically drawn. This arrow defines the polarity vector of a given cell (see Figure 2.a and 2.b). Figure 2.c shows the delineation result for all the cells of interest in our example dataset.

**Figure 2:**
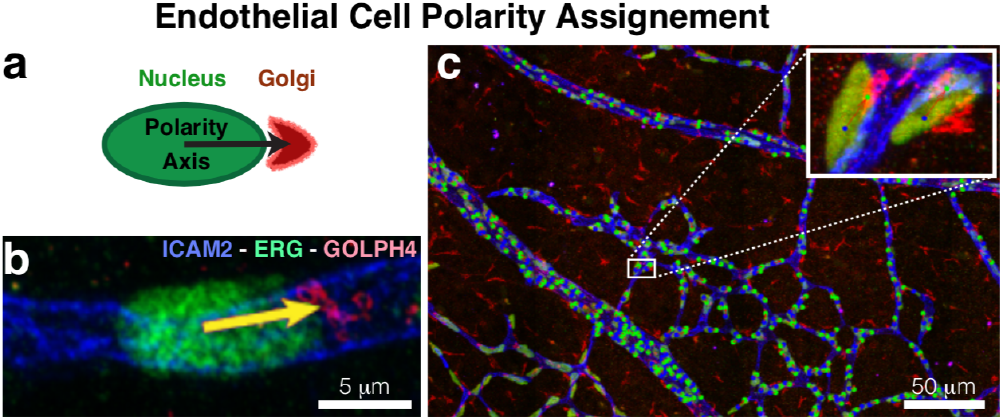
a) Polarity vectors are defined between the approximate centres of mass of the nucleus and the Golgi of any given cell. b) Example of polarity vector on Golph4-Erg-ICAM2 stained plexus. c) Plexus-wide view of the polarity vector delineation on Figure 1.a. Close-in panel provides a detailed view of the polarity delineation on a subset of endothelial cells.

Figure 3: presents the results of the PolNet analysis on the example dataset. Figure 3.a shows the overlay of the luminal mask and the cell polarity vectors in blue and the flow vectors (WSS selected on the dropdown menu) in red. Figure 3.b presents the quantification of the relative angles defined by each pair of polarity and flow vectors. It also includes the results of a statistical test for randomness in the distribution of these angles. Figure 3.c displays an example of subdivision into three regions (arterial, venous, and capillary) for further analysis. Finally, Figure 3.e shows the results of the analyses applied to each of the individually selected regions.

**Figure 3:**
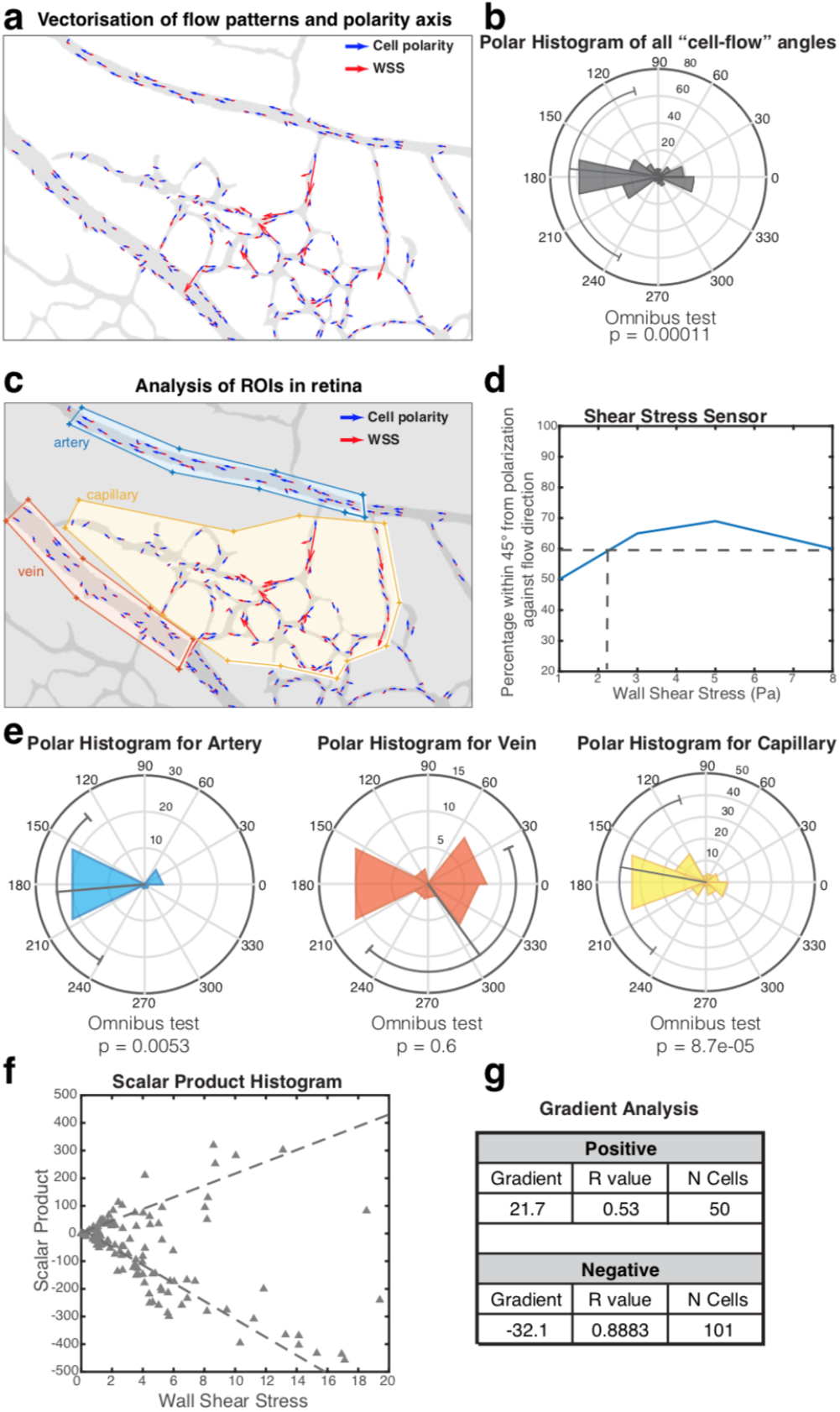
Polarity vs wall shear stress (WSS) analysis. a) Overlay of luminal mask (grey), polarity vectors (blue), and WSS vectors scaled according to magnitude (red). b) Polar histogram with relative angles formed by polarity and WSS vectors on each cell, results of statistical test for random distribution. c) Example of region subdivision: yellow, orange, and blue polygons classify the endothelial cells into arterial, capillar, and venous, respectively. d) Shear stress sensitivity analysis: percentage of cells polarised against the flow (within 45 degrees tolerance) as a function of WSS experienced. e) Results of the polarity analysis applied to each delineated region separately. f) Scatter plot showing the scalar product value for every pair of WSS and polarity vector. g) Scalar product quantification for both the positive and negative subgroups: gradient of the linear regression, Pearsons r-value, and number of cells.

## 4 Discussion

PolNet was conceived to be an easily extensible tool for network-level quantitative research in vascular morphogenesis. To date, PolNet has proven invaluable for the study of endothelial cell polarisation and migration during vascular patterning, as demonstrated by our recent papers [1, 2]. We have taken special care to make the pipeline easily deployable across a broad range of computer configurations for easy adoption of the Pol-Net software by the scientific community. The workflow described in this paper is optimised for the quantitative analysis of the relationship between endothelial cell polarity and haemodynamics in the neonatal mouse retina. However, the software can be easily extended to study the relationship between blood flow and other cellular/molecular processes relevant to vascular morphogenesis, including, but not restricted to, gene expression patterns, changes in endothelial cell morphology, proliferation and apoptosis rates, changes in vessel calibre or recruitment of mural cells. The only requirement is the ability to image and quantify the signal under investigation and compare it with the spatial distribution of flow parameters computed with the HemeLB CFD package. The current approach was developed for one of the most widely used animal models of angiogenesis, the mouse retina. However, other tissues of interest have more complex vascular architectures involving complex 3D vessel configurations. In the future, we plan to extend PolNet to include capabilities to segment and simulate haemodynamics in tissues and organs displaying a highly three-dimensional vascular organisation. With the advent of improved clearing techniques [19] and new imaging techniques [20], PolNet will be a powerful analysis method to address the complexity of endothelial cell biology at the network-level in intact organs.

## Author Contributions

Designed research: MOB, MLJ, PVC, HG, CAF; performed research: MOB, MLJ, RWN, AP, CAF; contributed analytic tools: RWN, AP; analyzed data: MOB, MLJ, AP, CAF; wrote the manuscript: MOB, MLJ, RWN, AP, PVC, HG, CAF.

## Acknowledgments

MLJ’s work was supported by the Francis Crick Institute which receives its core funding from Cancer Research UK (FC001999), the UK Medical Research Council (FC001999), and the Wellcome Trust (FC001999). HG is supported by an European Research Council (ERC) consolidator grant Reshape (311719), and by the British Council BIRAX initiative. CAF was supported by an ERC starting grant (679368), the H2020-Twinning grant (692322), the Fundação para a Ciência e a Tecnologia funding grants: IF/00412/2012; EXPL-BEX-BCM-2258-2013; PRECISE-LISBOA-01-0145-FEDER-016394; and LISBOA-01-0145-FEDER-007391, project cofunded by FEDER, through POR Lisboa 2020 - Programa Operacional Regional de Lisboa, PORTUGAL 2020, and Fundação para a Ciência e a Tecnologia. MOB acknowledges support from the UK Engineering and Physical Sciences Research Council under the project “UK Consortium on Mesoscale Engineering Sciences (UK-COMES)” (Grant No. EP/L00030X/1). This work used the ARCHER UK National Supercomputing Service (http://www.archer.ac.uk). The authors acknowledge the contributions of the HemeLB development team and the support of the UCL Research Software Development Group (RSD@UCL) in the completion of this work.

